# A Comprehensive Computational Model of the Human head for Designing and Optimizing Visual Brain-Machine Interfaces

**DOI:** 10.64898/2026.05.14.725091

**Authors:** Shengjian Lu, Tonghe Yang, Yuan Geng, Huan Wu, Yangrui Huang, Te Zheng, Haodi Chen, Shurui Huang, Yi Cao, Jian Yang, Wentao Yan, Yikui Zhang, Wencan Wu

**Author notes:** Corresponding author (Yikui Zhang); (Wencan Wu); (Wentao Yan); (Jian Yang). These authors contributed equally to this work (Shengjian Lu, Tonghe Yang).

## Abstract

Brain-machine interfaces (BMIs) for vision restoration require models that accurately simulate the anatomy and electrical properties of visual pathways. However, current models focus only on isolated structures, such as the retina or brain, and overlook surrounding tissues. Here, we present a comprehensive computational model of the human head, incorporating the entire visual pathway—including the eye, optic nerve, and brain—along with critical neighboring tissues such as the orbit, paranasal sinuses, enabling precise simulations. Validation using human and large animal data demonstrated a strong correlation between the simulated and measured electrical potentials. Component elimination analysis revealed that the optimized comprehensive model outperformed simplified versions. The model’s utility was demonstrated through multiple applications: (1) comparative analysis of electrical neuromodulation technologies for optic neuropathy, revealing the filed intensity limitations of noninvasive approaches and the safety concerns of invasive intraorbital approach; (2) identification of optimal stimulation site, revealing that transnasal stimulation at the optic chiasm outperformed traditional approaches; and (3) in silico design of electrode arrays for optic nerve prosthetics, demonstrating theoretical advantages in invasiveness and visual field coverage compared to existing retinal and cortical prosthetics. This validated and versatile computational resource supports the development of neuromodulation strategies and visual BMI technologies.

## 1. Introduction

Optic nerve repair represents a critical frontier in visual neuroscience. Optic nerve diseases—including glaucomatous, inflammatory, ischemic, and traumatic optic neuropathies—are the leading cause of irreversible blindness worldwide^1,2^. Recently launched initiatives, such as the Transplantation of Human Eye Allografts (THEA) program by the Advanced Research Projects Agency for Health (ARPA-H)^3^, have further heightened interest in technologies that enable optic nerve repair and eye-to-brain reconnection, with restoration of visual pathway function remaining the principal bottleneck to their success. While advances in biological strategies—for example, gene therapy and local delivery of neurotrophic factors—continue to drive progress in optic neuropathy management^4–7^, electrical neuromodulation via brain–machine interfaces (BMIs) offers complementary approaches. These BMIs can either promote guided axonal regeneration^8,9^ or deliver functional visual perception by replacing natural retinal signals^10,11^.

Despite growing interest in visual BMIs, most existing computational models focus on isolated anatomical structures^12–15^—such as the retina, optic nerve, or visual cortex—without incorporating the surrounding tissues that significantly influence electrical field propagation. This limitation hampers the accurate prediction of stimulation efficacy and safety, thereby slowing therapeutic development and clinical translation. Moreover, the lack of anatomically integrated models limits the ability to systematically evaluate and optimize emerging neuromodulation technologies. To address this gap, we developed a comprehensive computational resource that integrates the entire visual pathway together with neighboring tissues—including the orbit, paranasal sinuses, skull, cerebrospinal fluid, and muscle. The robustness of this model was validated by a strong correlation between simulated and recorded electrical potentials in both human and large animal studies. Systematic component elimination analysis further showed that the comprehensive model consistently outperformed simplified versions. Notably, the applications of our model include, but are not limited to, enabling robust assessment and optimization of electrical neuromodulation strategies for optic neuropathies informing the design of optic nerve prosthetic electrode arrays.

Together, these results position our validated computational platform as a robust and versatile tool for researchers to evaluate stimulation protocols, optimize electrode designs, and predict biological outcomes prior to costly in vivo experimentation. By making this resource accessible to the research community, we aim to accelerate the development of neuromodulation strategies and next-generation visual BMIs for blinding diseases.

## 2. Results

### 2.1 Personalized Head Model Reconstruction using Multimodal Neuroimaging

To achieve accurate segmentation of the visual pathway and surrounding structures for finite element modeling, we integrated cranial CT and dual T1-weighted MRI data from three young male adults (**Fig. 1A–B**). Both magnetization-prepared rapid gradient echo (MPRAGE) and magnetization-prepared 2 rapid acquisition gradient echo (MP2RAGE) MRI sequences were acquired to enable comprehensive visualization of cranial soft tissues and optimize contrast between gray and white matter, respectively^16,17^. Image registration was performed with the FMRIB Software Library (FSL) v6.0.4 to ensure precise spatial alignment of CT and MRI datasets (**Fig. 1C**).

**Figure 1.**
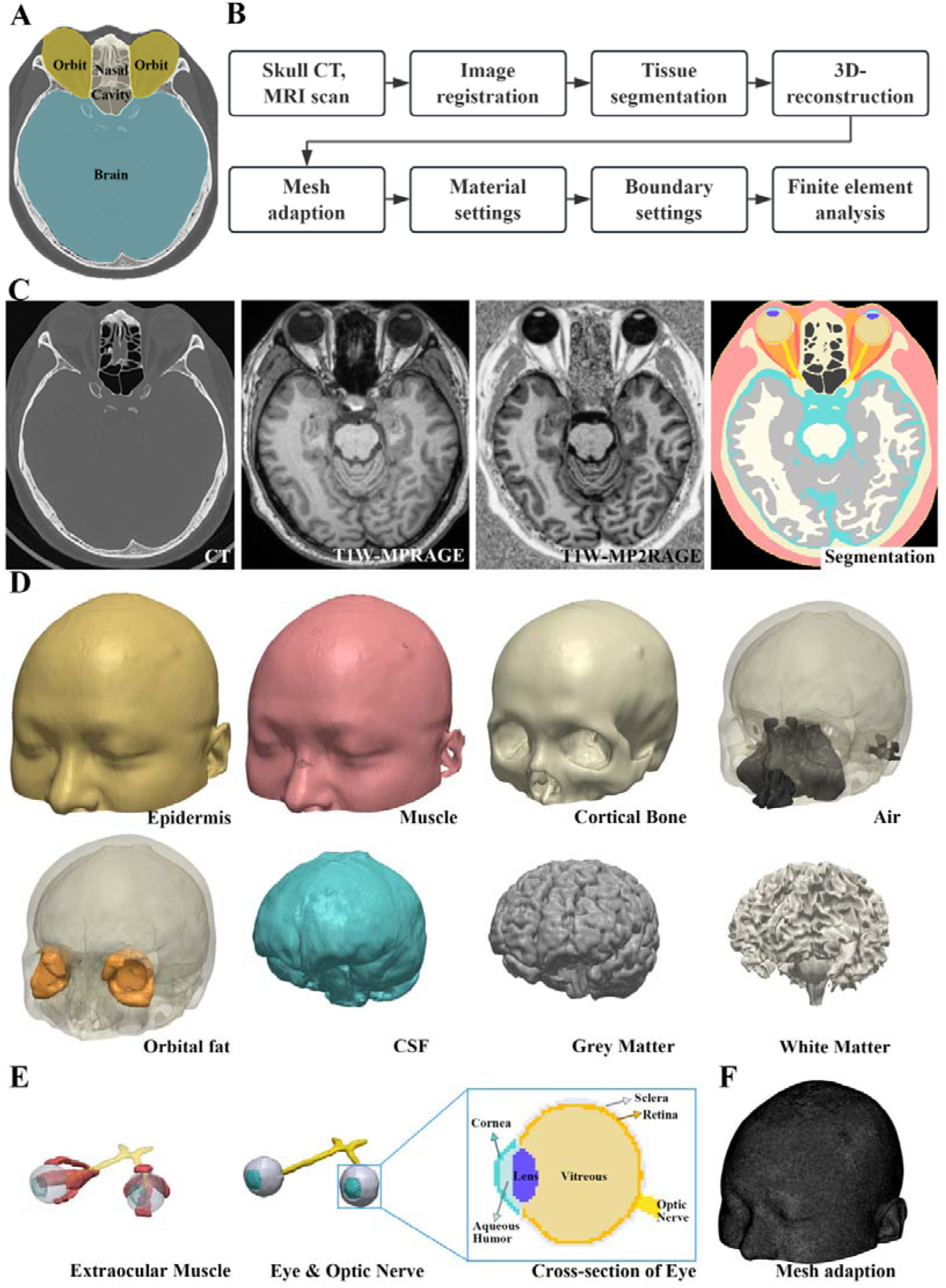
Modeling process of a personalized human head model. **(A)** Overview of the target regions including orbit, nasal cavity, and brain. **(B)** Flowchart of the modeling and finite element analysis workflow. **(C)** Demonstration of the image registration and tissue segmentation results. **(D)** 3D reconstruction of the anatomical structures, showing individual tissue components such as skin, muscle, cortical bone, air, orbital fat, cerebrospinal fluid (CSF), gray matter, and white matter. **(E)** 3D reconstruction of the extraocular muscles, the eyes (segmented into the cornea, aqueous humor, lens, vitreous humor, retina, and sclera), and the optic nerve. **(F)** Visualization of the adapted mesh applied to the entire skull.

Tissue segmentation was conducted using 3D Slicer (v5.6.2). We first delineated key structures from CT images, including low-signal air in the nasal cavities and paranasal sinuses, as well as high-signal cranial bones. Subsequent segmentation of cranial soft tissues was accomplished using MRI, permitting clear separation of gray and white matter within the cranial cavity. Intracranial interstitial spaces between neural tissues were systematically classified as cerebrospinal fluid (CSF) (**Fig. 1D**). Residual soft tissues identified by head contour were segmented and classified as muscle. The surfaces of muscular tissue were extended by 0.5 mm to define a skin layer, which is important for accurately modeling spatial variations in electrical parameters^18^. Ocular structures were then segmented in detail (**Fig. 1E**), including the anterior chamber, lens, and vitreous body. The surface of the anterior chamber was extended anteriorly by 0.5 mm to define the cornea, and both the retina and sclera were delineated as 0.5 mm layers surrounding the vitreous body. The optic nerve, extraocular muscles, and orbital adipose tissue were also systematically segmented.

Following tissue segmentation, we systematically addressed minor surface discontinuities and local geometric irregularities throughout the entire model, thereby facilitating more efficient finite element mesh generation and refining its overall computational quality **(Fig. 1F)**. These procedures enabled construction of a personalized, comprehensive computational model that integrates the entire visual pathway and all surrounding tissues for electrical simulation.

### 2.2 Validation of the Head Model through in Vivo Stimulation and Measurements

To rigorously validate the accuracy of our computational model, we performed noninvasive electrical stimulation experiments in human participants. A ring-shaped stimulating electrode was placed at the frontal pole midline (Fpz) and a reference electrode at the occipital region (Oz), delivering direct current stimulation at 1, 2, and 4 V. Electrical potentials were measured at four neuroanatomically defined locations: frontal midline (Fz), parietal midline (Pz), left temporal midpoint (T3), and right temporal midpoint (T4). Simultaneously, we simulated identical stimulation parameters in our head model and extracted corresponding electric potential values from these sites.

Comparative analysis revealed a robust correlation between measured cranial potential distributions and simulated results, with correlation coefficients improved with increasing stimulation intensity (**Fig. S1**). To comprehensively assess simulation fidelity, we systematically investigated the impact of tissue component modifications, including addition of cancellous bone, and elimination of the lens, aqueous humor, air cavities, orbital adipose, CSF, or muscular tissue (**Fig. S1A-F**). Notably, alterations to individual tissue components had minimal effect on simulation accuracy (**Fig. S1G**), likely due to the superficial positioning of recording electrodes, where the electric field is dominated by proximal anatomical structures rather than deeper tissues.

Although human validation demonstrated strong correlations, measurements were limited to superficial recording sites due to ethical constraints. To further validate our model at deeper anatomical sites, we employed a large animal (goat) model, which enabled microinvasive placement of electrodes at the optic nerve^19^. A ring-shaped stimulating electrode was positioned on the cornea, and a half-ring reference electrode was placed at the optic chiasm via transnasal endoscopy (**Fig. 2A–C**), generating an electric field along the optic nerve relevant to directional regeneration. Recording electrodes were inserted into the bilateral retrobulbar and pre-chiasmatic optic nerves, as well as at Oz. After 2 V direct current stimulation, potentials were recorded via these implanted electrodes. Postoperative CT and MRI were taken for detailed anatomical reconstruction and precise simulation (**Fig. 2D–F**). Simulation results demonstrated a gradient potential along the retrobulbar optic nerve toward the optic chiasm as expected (**Fig. 2G**), with a strong correlation (r = 0.9378) between in vivo measurements and computational predictions (**Fig. 2H**).

**Figure 2.**
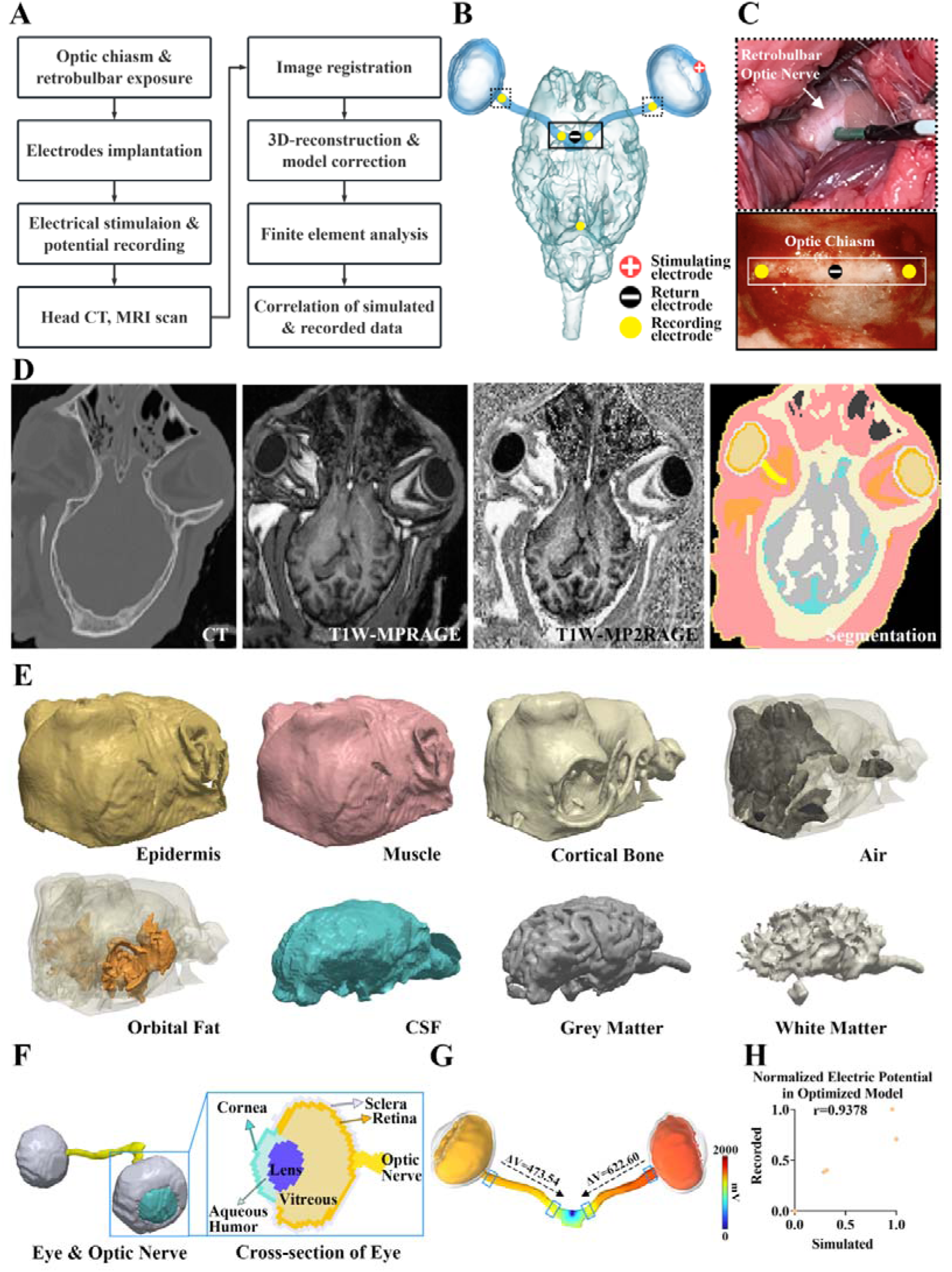
Measurements and validations of the electric potential distribution in simulation models for goats. **(A)** Flowchart outlining the measurement and finite element analysis processes conducted on goats. **(B)** Schematic illustration showing the positions of the electrodes in a goat. **(C)** Implantation of the recording electrode at the retrobulbar optic nerve and optic chiasm. **(D)** Demonstration of the image registration and tissue segmentation results. **(E)** 3D reconstruction of various anatomical structures, including skin, muscle, cortical bone, air cavities, orbital fat, CSF, gray matter and white matter. **(F)** Structural model of the eye and optic nerve, with an enlarged cross-sectional view showing internal components (cornea, lens, aqueous humor, vitreous, retina, sclera, and optic nerve). **(G)** Illustrations of the electric potential distributions in the retina, optic nerve and brain. **(H)** Correlation between the normalized recorded and simulated electric potentials (r=0.9378).

### 2.3 Validation of the Head Model through systematic elimination of tissue components

Systematic elimination of tissue components in the large animal model with deep electrode recordings showed that omitting structures such as nasal air, orbital fat, CSF, and skull muscle each reduced the correlation coefficient from r = 0.9378 to r = 0.9204, 0.9188, 0.9141, and 0.7683, respectively (**Fig. S2**). These results underscore that a comprehensive model including essential surrounding tissues is necessary for reliable electric field simulations. Interestingly, incorporation of cancellous bone slightly reduced simulation accuracy (r = 0.9292) (**Fig. S2A**), consistent with prior research^14,20^.

### 2.4 Non-invasive Therapeutic Neuromodulation Technologies: Low Electrical Field Intensity Along the Optic Nerve

We first used our model to evaluate the efficiency of current noninvasive neuromodulation techniques^21–26^. Prior work indicates that applied electric fields may promote axonal regeneration^26–29^, but clinical benefits in optic neuropathies have been inconsistent, potentially due to insufficient electric field strength delivered to the optic nerve.

Using our validated head model, we analyzed electric field distributions along the optic nerve in three human subjects under four commonly used noninvasive stimulation protocols. In the frontal-occipital protocol (Fig. 3A), with electrodes positioned along the skull midline, the potential difference across both optic nerves was negligible (0.06 mV; Fig. 3B–C). The orbital-occipital (Fig. 3D) and corneal-occipital (Fig. 3G) protocols produced higher ipsilateral potential differences (0.46 mV and 0.81 mV, respectively; Fig. 3E–F, H–I), while maintaining minimal contralateral effects. The corneal concentric ring protocol, in which concentric electrodes were placed directly on the cornea (Fig. 3J), generated the largest ipsilateral potential difference (2.43 mV), with a low contralateral effect (−0.15 mV; Fig. 3K–L). All these noninvasive protocols produced electric field intensities across the optic nerve (length: 40 mm) at least three orders of magnitude below the threshold required for effective directional axonal regeneration as established by both in vivo and in vitro studies (100 mV/mm)^9,27^.

**Figure 3.**
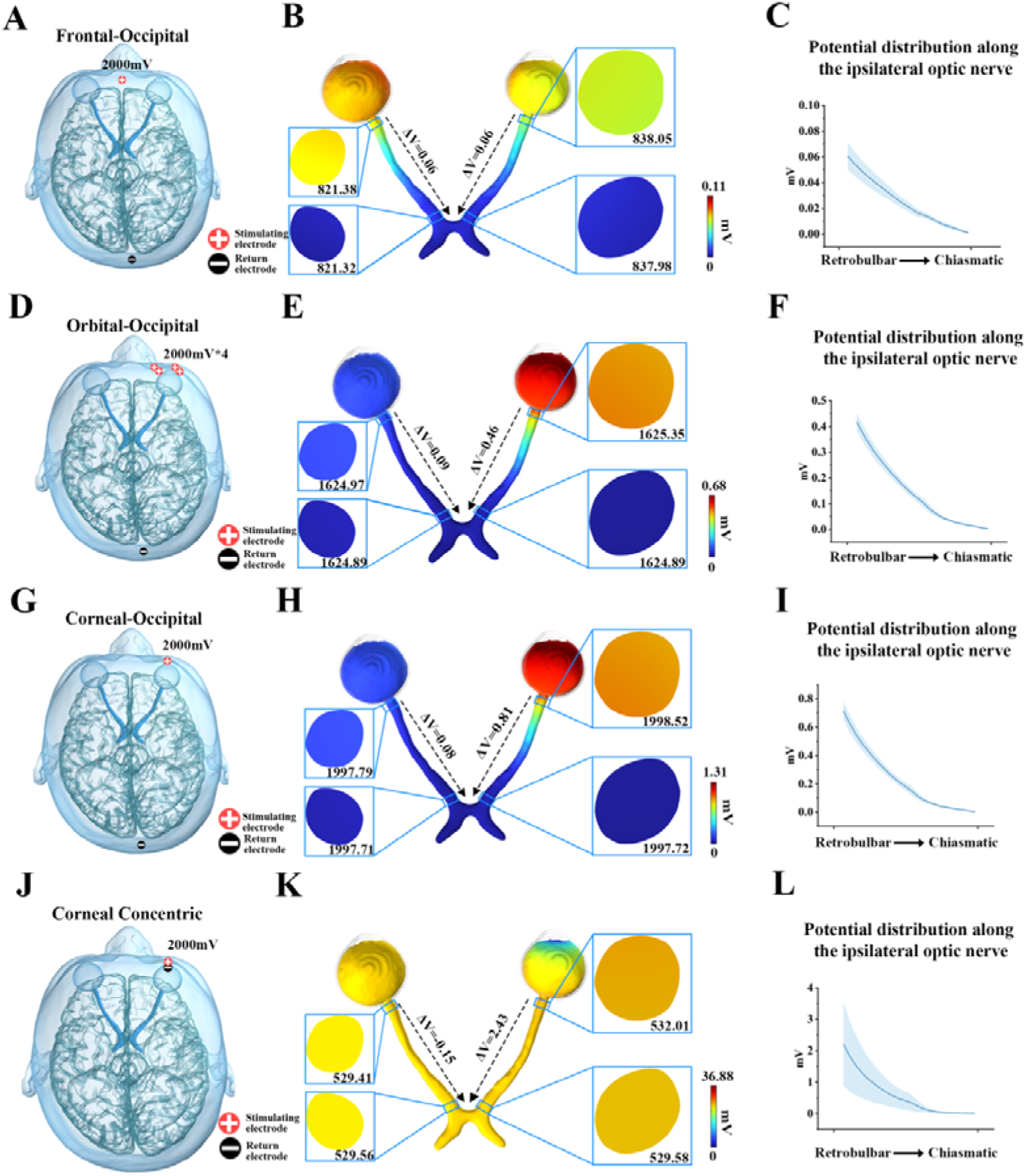
Distribution of electrical potential in the pre-chiasmatic visual pathway under noninvasive stimulating approaches. **(A, D, G, J)** Electrode arrangements under the frontal-occipital, orbital-occipital, corneal-occipital, and corneal concentric rings protocols, respectively; **(B, E, H, K)** Illustrations of the electric potential distribution along the optic nerve corresponding to each protocol; **(C, F, I, L)** Quantifications of the electric potential distribution along the optic nerve corresponding to each protocol, n = 3, data are presented as means ± s.e.m.s.

### 2.5 Invasive Therapeutic Neuromodulation Technologies: Higher Electrical Field Intensity with Safety Concerns

We next evaluated two intraorbital cuff electrode protocols, which require open-orbit surgery for placement on the optic nerve^30^. The corneal-orbital apex protocol (Fig. 4A) generated a large potential difference (ΔV = 1272 mV) along the ipsilateral optic nerve (Fig. 4B–C), several hundred times higher than noninvasive methods. The retrobulbar-orbital apex protocol (Fig. 4D) produced an even greater potential difference (ΔV = 1443 mV) and reduced contralateral optic nerve stimulation from 21 mV to 11 mV (Fig. 4E–F), indicating greater efficacy and selectivity (Fig. 4G). However, in rat experiments, intraorbital cuff electrode implantation led to substantial loss of RBPMS-positive retinal ganglion cells (RGCs) at two weeks post-implantation (Fig. 4H–K), likely due to mechanical compression from the electrode.

**Figure 4.**
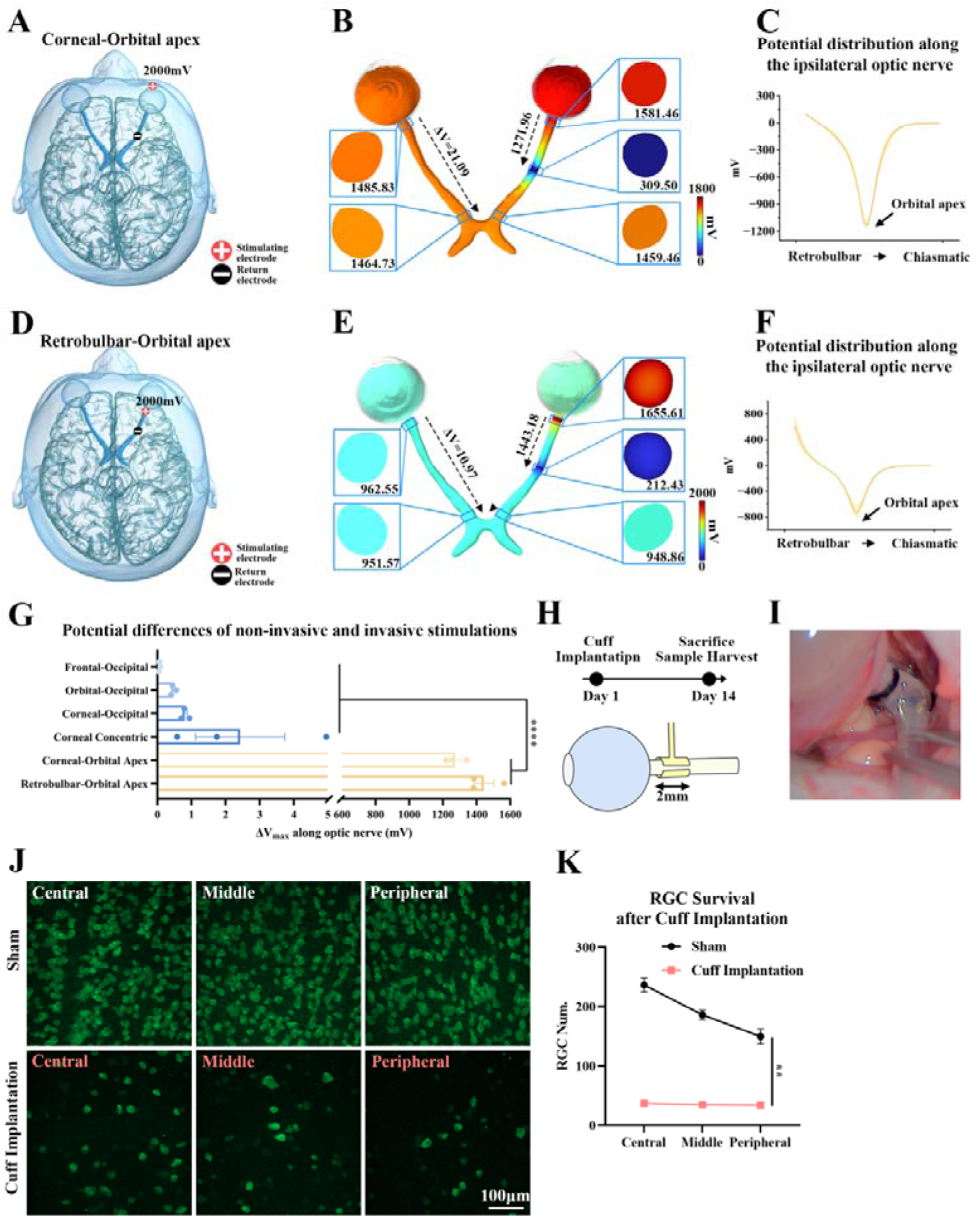
Advantages and safety issues of intraorbital stimulation approaches. **(A, D)** Electrode arrangements under the corneal-orbital apex and retrobulbar-orbital apex protocol; **(B, E)** Illustrations of the electric potential distribution along the optic nerve corresponding to each protocol; **(C, F)** Quantifications of the electric potential distribution along the optic nerve corresponding to each protocol, n = 3, data are presented as means ± s.e.m.s.; **(G)** Comparison of the maximum potential difference (ΔV) along the optic nerve across various noninvasive and intraorbital stimulation strategies. Two-Way Anova, n = 3, data are presented as means ± s.e.ms, ****: p < 0.0001. **(H)** Experimental design for the safety evaluation of the cuff electrode implantation process. **(I)** Microscopic view of the implanted cuff on the retrobulbar optic nerve. **(J)** Immunofluorescence staining of RBPMS-positive retinal ganglion cells (RGCs) in the central, middle, and peripheral retina in the sham and cuff-implanted groups. **(K)** Quantitative analysis of RGCs in all retinal zones following cuff electrode implantation compared with that in the sham group. Two-Way Anova, n = 3, data are presented as means ± s.e.ms, **: p < 0.01.

### 2.6 Identification of Optimal Stimulation Site Using a Transnasal Approach

To address the limitations of conventional methods, we leveraged a transnasal endoscopic approach, a well-established technique in modern neurosurgery^31,32^. This approach provides safe and direct access for implantation of the BMI at the optic canal or the anterior optic chiasm. Unlike the orbital segment of the optic nerve, the intracanalicular optic nerve and the optic chiasm do not move during eye movements. Furthermore, the air-filled sphenoid sinus provides a spacious room for BMI implantation.

We proposed placing a 180° half-ring electrode on the anterior optic chiasm during the surgery, paired with a corneal electrode (Fig. 5A–B). Simulations demonstrated a substantial potential difference along the ipsilateral optic nerve (ΔV = 739.4 mV; Fig. 5C–D), comparable to that of traditional intraorbital electrodes (Fig. 5E). Notably, the corneal-chiasmatic approach (2 V stimulation) induced a maximum brain electric field intensity of 0.07 mV/mm at the skull base and chiasm (Fig. S3), which is well below the established safety threshold for brain tissue (0.4–0.8 mV/mm)^33,34^, indicating a favorable safety profile for clinical application.

**Figure 5.**
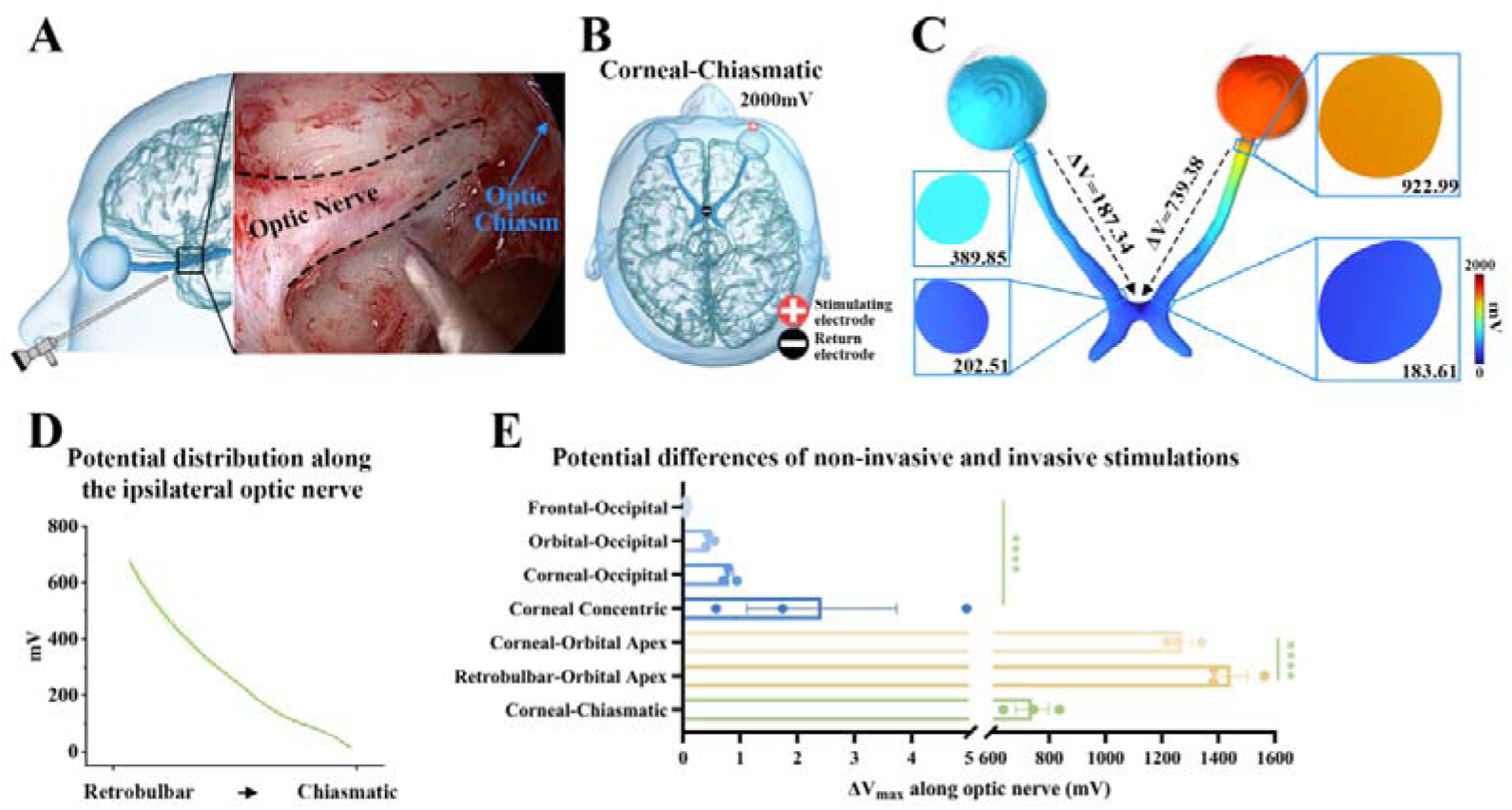
Identification of transnasal electrical stimulation at the optic chiasm outperformed traditional approaches. **(A)** Illustration and endoscopic view of the endoscopic endonasal approach for exposing the optic chiasm. (**B)** Electrode arrangement used in the corneal-chiasmatic protocol. **(C)** Simulated electric potential distribution along the optic nerve. **(D)** Quantification of the potential distribution along the ipsilateral optic nerve, n = 3, data are presented as mean ± s.e.m. **(E)** Comparative analysis of the maximum potential difference (ΔV) along the optic nerve across various stimulation protocols. Two-Way Anova, n = 3, data are presented as mean ± s.e.m, ****: p < 0.0001.

### 2.7 Proof-of-Concept Designs for Optic Nerve Prosthetic Arrays

Another application of our computational model was to investigate electrode designs for optic nerve BMIs implanted at the nerve surface, offering a potential alternative to retinal or cortical visual prosthetics. A key question is whether a surface-mounted BMI can activate axons located deeper within the nerve. By varying electrode size, channel count, spatial arrangement, and return electrode position, we found that both the strength and direction of the electric field within the optic nerve could be modulated, suggesting that with appropriate design, any region of the optic nerve could theoretically be activated.

Comparisons between 0.5 mm and 1 mm electrodes placed on the optic nerve surface indicated that smaller electrodes produced more concentrated electric fields with greater intensity along the radial axis, whereas larger electrodes generated broader, but lower-intensity, field distributions (Fig. 6A–D). Increasing the number of active channels from 4 to 16 in electrode arrays reduced the electric field intensity along the radial axis (Fig. 6E–H), likely due to a rise in central nerve potential (Fig. S4).

**Figure 6.**
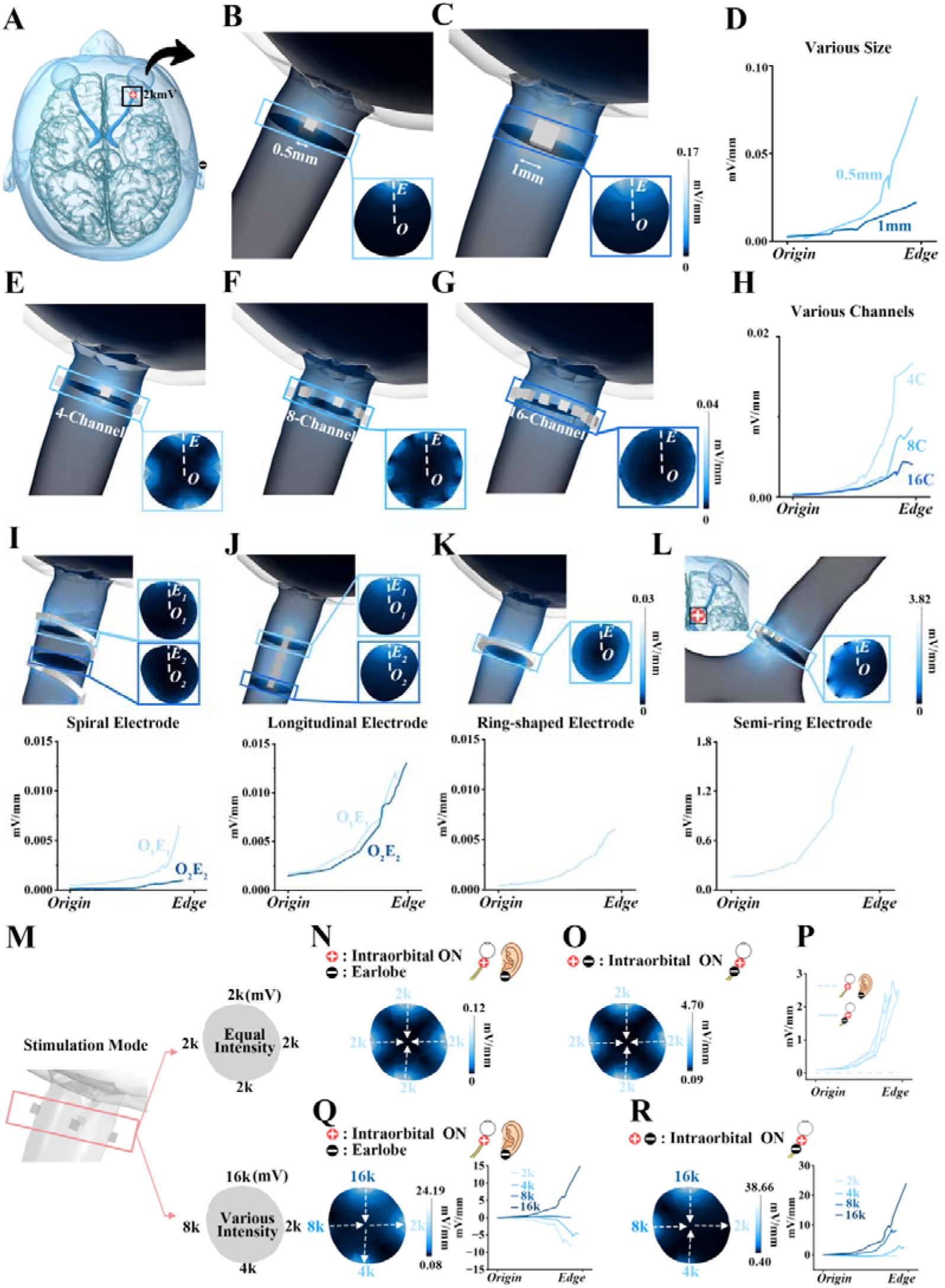
Demonstrations of various electrode designs for optic nerve prosthetics. **(A)** Position of the optic nerve prosthesis. **(B, C)** Schematics of 0.5 and 1 mm electrode configurations and the cross-sections of the electrode sites. **(D)** Quantification of the electric field on the cross-section generated by electrodes of different sizes. **(E-G)** Schematics of 4 to 16-channel electrode configurations and the cross-sections. **(H)** Quantification of the electric field on the cross-section generated by different channel configurations. **(I-L)** Schematics and quantifications of the electric field distributions on the cross-section generated by spiral electrodes, longitudinal electrodes, ring-shaped and semi-ring electrodes. **(M)** Illustration of equal stimulation intensity and varying stimulation intensities among the channels in the 4-channel setup. **(N-P)** Electric distributions on the cross-section under stimulation with equal intensity (2 V) and return electrodes placed on the earlobe and intraoribital optic nerve. The arrows indicated the directions of the electric field. **(Q, R)** Electric distributions on the cross-section under stimulation with various intensities (2-16 V), with the return electrodes placed on the earlobe and intraoribital optic nerve. The arrows indicated the directions of the electric field.

Exploring different spatial configurations—including spiral, longitudinal, ring-shaped, and semi-ring designs—revealed that each arrangement produced distinct internal electric field patterns within the optic nerve (Fig. 6I–L). With equal voltage inputs, positioning the return electrode at the intraorbital optic nerve resulted in a much larger electric field compared to placement on the earlobe (Fig. 6M–P). Furthermore, varying voltage input across channels and adjusting the return electrode location allowed control over both the direction and the intensity of electric fields within the optic nerve. For instance, when the return electrode was on the earlobe, electric fields near the lower voltage electrodes (2 and 4 V) were directed toward these electrodes (Fig. 6Q), while positioning the return electrode at the intraorbital optic nerve generated electric fields directed away from all electrodes except the 2 V channel.

## 3. Discussion

In this study, we established the first high-resolution, subject-specific computational model integrating the entire visual pathway—from the cornea through the optic nerve, orbit, nasal structures, and brain. By combining CT and MRI imaging, we achieved detailed segmentation of both soft and hard tissues, enabling accurate simulation of electrical field distributions throughout these anatomically connected regions. Importantly, the model’s predictions were validated by strong correlations with in vivo electrical measurements—both in human patients (superficial recordings) and in large animal models (deep optic nerve recordings)—demonstrating its reliability and relevance for translational applications. Using this validated model, we showcased its utility through three key applications: (1) comparing neuromodulation strategies for optic neuropathy, highlighting limitations of noninvasive and safety issues of invasive methods; (2) identifying transnasal optic chiasm stimulation as an optimal and safer site compared to traditional approaches; and (3) designing and evaluating optic nerve prosthetic arrays. This resource addresses a major gap in previous modeling efforts and offers new insights into designing and optimizing next-generation neural stimulation strategies and visual prostheses.

### Comparing with Previous Computational Models

By leveraging detailed computational modeling and finite element analysis, previous studies have made significant advances in the field; however, these works did not comprehensively include the entire visual pathway along with the surrounding orbit and nasal structures. For instance, Xiaofan Su et al. developed a high-resolution, full-eye model—including the cornea, aqueous humor, lens, vitreous body, retina, choroid, and sclera—to simulate retinal electrical stimulation with spatial selectivity. However, their model did not incorporate the optic nerve and brain^12^. Sangjun Lee et al. extended retinal stimulation models by incorporating the skull and brain, allowing for a more comprehensive investigation of how transcranial electrical stimulation affects the retina; however, they excluded the optic nerve and the surrounding orbital and nasal structures^13^. Similarly, Huang et al. constructed anatomically detailed human head models to analyze transcranial electrical stimulation, but did not incorporate the eye, optic nerve, orbit, and nasal cavity^14^. In contrast, our study established a comprehensive model and verified its importance through systematic elimination of tissue components in the large animal model: for example, omitting nasal air or orbital fat in the model significantly reduced the correlation coefficient between the simulated and measured electrical distribution (**Fig. S2**).

Additionally, previous head models typically relied solely on MRI^35,36^, which provides limited contrast for the skull, CSF, and air cavities—tissues with vastly different conductivities—resulting in challenges for accurate segmentation. By integrating both CT and MRI data, our pipeline achieves precise tissue segmentation, enabling accurate reconstruction of tissue boundaries that are critical for realistic electrical field simulations. Furthermore, the use of a large animal model provided distinct advantages for validating simulated electrical distributions. Unlike human studies, the large animal model allowed direct electrode insertion into the optic nerve and visual pathway, enabling precise measurement of electrical fields. Additionally, immediate post-operative CT and MRI imaging were available to capture anatomical changes following electrode implantation via craniotomy—changes crucial for accurate modeling. In contrast, this immediate post-operative imaging is rarely available in human studies. Incorporating these real-time anatomical updates significantly enhanced the reliability of our model validation and its correlation with measured electrical data.

### Optic Nerve BMI: A Promising Pathway Beyond Traditional Visual Prosthetics

Optic nerve BMI represents a promising alternative to current visual prosthetic technologies. The surface areas of the retina (962_–_1857 mm²)^37^ and the primary visual cortex (2938–6296 mm²)^38^ are substantially larger than the intraorbital optic nerve’s cross-sectional area (3.8-8 mm²)^39^ in human adults. Consequently, existing retinal or cortical prosthetics face limitations such as restricted visual field coverage and invasiveness associated with implantation via retinal detachment or open-skull surgery ^40–43^. The larger the visual field coverage desired, the larger—and more invasive—the BMI system typically becomes.

In contrast, an optic nerve BMI could theoretically activate a greater number of axons, enabling a broader visual field with a much smaller implant size^10,11,44–46^, potentially implanted via minimally invasive trans-nasal endoscopy^19^. However, significant technical challenges remain. These include the need for ultra-high spatial and temporal precision in electrical field distribution across the optic nerve to selectively target individual axons. Furthermore, the dura mater and other meningeal layers may impede effective signal transmission, necessitating further research into optimized electrode designs and stimulation parameters.

### Necessity of personalized model

Personalized models are necessary for accurately capturing the subtle yet crucial individual anatomical differences that potentially influence simulation outcomes^36,47^. In this study, although the three subjects showed similar overall electrical field distributions, their electric potentials varied especially under corneal concentric ring stimulation (**Fig. 3L**). Huang et al. also demonstrated that ignoring personal anatomic differences often leads to inaccuracies in predicting neural responses^14^. However, the development of a personalized comprehensive computational model remains technically challenging and time-consuming, which took 3-5 days in this study. Advances in artificial intelligence could potentially streamline this process and improve modeling accuracy.

### Limitations

Our study has several limitations. First, the anatomical accuracy was constrained by MRI resolution (1 mm), which necessitated exclusion of sub-millimeter structures such as the meninges and subcutaneous tissue. Manual segmentation also introduced variability, especially for complex structures like the irregular skull, paranasal sinuses, and CSF spaces. Additionally, the lack of precise conductivity values for direct current stimulation required us to use 20 Hz alternating current conductivity data instead. Although Huang et al.14 reported minimal conductivity variation within the 100 Hz range^14^, this substitution remains a methodological consideration.

Furthermore, our model represents the optic nerve as a uniform cylinder without the detailed anatomical features of actual nerve tissue, such as glial cells, blood vessels, and extracellular matrix. Future studies may benefit from 3D reconstruction of the optic nerve using electron microscopy to improve anatomical fidelity. Finally, current electrode array designs do not adequately account for the temporal dynamics of electrical field changes. Future research should address these limitations and focus on developing strategies for the selective activation of targeted axon bundles in optic nerve prosthetics.

## 4. Methods

### Study Design

In this study, we aimed to develop a finite element model of the orbital-nasal-cerebral region on the basis of skull imaging data obtained from volunteers to investigate the efficacy and safety of different optic nerve stimulation protocols. Three adult male volunteers were recruited, and CT and MRI scans were obtained to construct personalized head models. In vivo measurements of electrical potential were taken from volunteers and a goat to validate the accuracy of the models. Using these models, we simulated seven electrical stimulation protocols, each characterized by a unique arrangement of electrodes or implantation setups, as well as various electrode designs for optic nerve prosthetics. Experiments on rats (n = 3) were conducted to assess the safety of intraorbital electrodes. Ethical approval was granted by the Ethics Committee of the Eye Hospital of Wenzhou Medical University (Approval No. 2024-204-K-171-01), and informed consent was obtained from each volunteer. The animal protocols were approved by the Experimental Animal Ethics Committee of Wenzhou Medical University (Approval No. wydw2024-0241).

### Animals

Male Sprague–Dawley (SD) rats were purchased from Beijing Vital River Laboratory Animal Technology Co., Ltd., and housed in the animal facility under temperature-and humidity-controlled conditions with a 12-hour light dark cycle. Rodent chow and water were provided ad libitum. All the rats were habituated for at least one week before the experiments commenced, and the rats were aged 2–3 months at the beginning of the study. Male saanen goats aged 4 to 7 months, with body weights of 20–25 kilograms, were purchased from the Caimu Livestock Company (Hangzhou, China) and housed in the animal facility at Wenzhou Medical University. They were kept in an air-conditioned room with a normal light/dark cycle and had free access to food and water. The goats also underwent a habituation period of at least one week before the beginning of the experiments.

### Image Acquisition and Registration

CT and MR imaging data were acquired to ensure comprehensive anatomical coverage and enable multimodal analysis. CT imaging was performed on a Siemens SOMATOM go. CT scanner (slice thickness = 0.6 mm, X-ray tube current= 146 mA, Hr60). MRI (Siemens Lumina) data were collected using two T1-weighted sequences: (1) a three-dimensional (3D) magnetization-prepared rapid gradient echo (MPRAGE) sequence (echo time (TE) = 3.41 ms, repetition time (TR) = 2200 ms, inversion time (TI) = 926 ms, slice thickness = 0.94 mm) and (2) a 3D magnetization-prepared 2 rapid acquisition gradient echo (MP2RAGE) sequence (TE = 2.89 ms, TR = 5000 ms, slice thickness = 1.14 mm)

Image registration was performed using the FMRIB Software Library (FSL) v6.0.4 to precisely align the CT and MR images. Initially, the CT image was registered to the MR image acquired with the MPRAGE sequence using the FLIRT tool with six degrees of freedom (DOFs) and a mutual information cost function, generating a transformation matrix and an aligned CT image. The MR images acquired with the MPRAGE sequence were subsequently aligned to the MR images acquired with the MP2RAGE sequence using FLIRT with 12 DOF and the same cost function, producing a second transformation matrix. The two matrices were then concatenated using the convert_xfm tool to create a combined transformation matrix, which was applied to the CT image to generate the final aligned CT image in the MRI space of the MP2RAGE sequence. This workflow, which is based on mutual information and affine transformations with trilinear interpolation, minimizes distortion and ensures high anatomical consistency, providing a robust foundation for subsequent analyses such as tissue segmentation, structural quantification, and image fusion.

### Tissue Segmentation and 3D Modeling

The registered CT and T1-weighted image data were imported into 3D Slicer (v5.6.2) for analysis, where manual segmentation of the following tissues was performed on the basis of different grayscale values: skin, muscle, cortical bone, cancellous bone, air cavities, CSF, gray matter, white matter, orbital fat, extraocular muscles, optic nerve, and ocular components (including the cornea, sclera, aqueous humor, lens, vitreous humor, and retina).

Owing to limitations in imaging resolution, very small structures, such as blood vessels, the dura mater, the pia mater, and nerve sheaths, were not segmented. Additionally, other cranial nerves were excluded from the segmentation process because they fell outside the primary focus of this study. For particularly thin tissues that exhibited poor contrast, such as the bony wall, retina, and cornea, a meticulous manual editing process was performed to correct incomplete boundaries, ensuring geometrically coherent representations of key tissue regions. After segmentation, any minor holes or irregularities were manually corrected. All manual segmentation procedures were carried out by an experienced PhD student in medicine.

### Electrical Stimulation Protocol and Electrode Modeling

The following seven electrical stimulation protocols were designed.

Noninvasive approaches: (1) frontal-occipital protocol: the stimulating electrode was positioned at Fpz, with the return electrode at Oz; (2) oribital-occipital protocol: the stimulating electrodes were evenly arranged around the periorbital region (four in total), with the return electrode at Oz; (3) corneal-occipital protocol: the stimulating electrode was positioned at cornea, with the return electrode at Oz; (4) corneal concentric rings protocol: an inner ring with a 5 mm diameter served as the stimulating electrode, and an outer ring with a 10 mm diameter served as the return electrode.

Intraorbital invasive approaches: (1) corneal-orbital apex protocol: the stimulating electrode was positioned at cornea, with the return electrode at orbital apex; (2) retrobulbar-orbital apex protocol: the stimulating electrode was positioned at retrobulbar optic nerve, with the return electrode at orbital apex.

Microinvasive approach: corneal-chiasmatic protocol: the stimulating electrode was positioned at cornea, with the return electrode at chiasm.

Noninvasive corneal/skin electrodes were constructed from copper rings with a diameter of 5 mm and a thickness of 0.5 mm. For all noninvasive protocols, conductive paste was applied between the skin and electrodes to ensure good electrical contact. The retrobulbar and orbital apex electrodes were also copper rings, with diameters ranging from 4 to 6 mm, whereas the chiasmatic electrode was a 180° half-ring. Each electrode was placed in the model and geometrically fitted to the curved surfaces as needed.

### Mesh Generation and Finite Element Simulation

The combined model (head and electrodes) was imported into COMSOL Multiphysics 6.2. The mesh generation strategy was optimized to account for the model’s anatomical complexity and the available computational resources. Tetrahedral elements were used to discretize the entire domain, with finer local meshes applied in thin or geometrically complex regions (e.g., cornea, retina) to capture high-gradient fields. Mesh convergence tests were performed to balance accuracy and computational efficiency.

The infinity analysis was executed using the stationary study of the AC/DC module. Tissue electrical properties (conductivity and relative permittivity) at 20 Hz were assigned to brain tissues, the skull, ocular components, and other relevant tissues. The electrodes were modeled as copper, with a relative permittivity of 1 and conductivity of 5.96×10^7^ S/m. The specific tissue electrical property values are provided in Table 1. The stimulating electrode was set to 2 V, the return electrode was set to 0 V, and all external surfaces were assigned electrical insulation conditions. The resistance of the reference electrode was set to 50 ohms by default.

**Table1:** Tissue electrical properties.

| Tissue | Conductivity (S/m) | Relative permittivity |
| --- | --- | --- |
| Air | $1 \times 10^{-14}$ | 1 |
| Skin | $2 \times 10^{-4}$ | $1.14 \times 10^3$ |
| Muscle | $2.07 \times 10^{-1}$ | $2.43 \times 10^7$ |
| Cortical bone | $2.00 \times 10^{-2}$ | $2.51 \times 10^4$ |
| Sponge bone | $7.89 \times 10^{-2}$ | $4.02 \times 10^6$ |
| grey matter | $4.31 \times 10^{-2}$ | $3.14 \times 10^7$ |
| White matter | $3.89 \times 10^{-2}$ | $1.75 \times 10^7$ |
| CSF | 2 | 109 |
| Orbital fat | $3.94 \times 10^{-2}$ | $2.09 \times 10^6$ |
| Optic nerve | $2.38 \times 10^{-2}$ | $8.07 \times 10^6$ |
| Retina | $4.31 \times 10^{-2}$ | $3.14 \times 10^7$ |
| Sclera | $5.02 \times 10^{-1}$ | $1.10 \times 10^6$ |
| Cornea | $4.18 \times 10^{-1}$ | $8.10 \times 10^6$ |
| Aqueous fluid | 2 | $1.09 \times 10^2$ |
| Vitreous | 1.5 | 99 |
| Lens | 0.2 | $2.13 \times 10^3$ |
| Conductive paste | 1 | 100 |

Under the quasistatic approximation of Maxwell’s equations, the following relationships were considered:

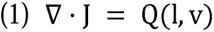

For passive tissues, Q(l, v) = 0 to ensure charge conservation.

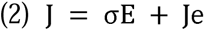

J includes the tissue conduction current (σE) and the externally injected current (Je).

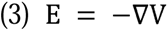

The electric field E is determined by the gradient of the electric potential V. Once V is determined, E and J can be computed with the model.

The finite element method was applied to numerically resolve these complex electric field equations, utilizing a robust edge element discretization strategy complemented by advanced preconditioning algorithms to efficiently resolve the sparse linear systems. After these equations were solved, 3D distributions of the electric potential (V), electric field (E), and current density (J) were obtained. The magnitude and distribution of the electric fields in key regions (e.g., optic nerve, chiasm, and visual cortex) were analyzed, enabling comparisons of the electric fields generated with different electrode placement strategies.

### In Vivo Electrical Stimulation and Measurements in Humans and Goats

In the human study, noninvasive electrical stimulation was provided using a ring-shaped stimulating electrode placed at Fpz and a reference electrode placed at Oz. Direct current stimulation of 1 V, 2 V, and 4 V were delivered, with electrical potential measurements recorded from four neuroanatomically defined locations: Fz, Pz, T3, and T4.

For the goat study, microinvasive electrical stimulation was provided by positioning a ring-shaped stimulating electrode on the cornea and a half-ring reference electrode at the optic chiasm. Recording electrodes were implanted in key locations, including the bilateral retrobulbar optic nerves, bilateral pre-chiasmatic optic nerves, and occipital region (Oz). After delivering 2 V of direct current stimulation, potential values were recorded from these electrodes.

### Optic Chiasm Transsphenoidal Endoscopic Exposure in Goats

The procedure followed a previously published protocol. Anesthesia was induced using intravenous propofol (10 mg/kg) and maintained with 3% isoflurane mixed with oxygen and air at a flow rate of 2 L/min (RWD Life Science Co., Ltd., China), which was delivered via a mechanical ventilator. To minimize intraoperative bleeding, hemocoagulase atrox (2 units per goat) was administered. Prior to surgery, the surgical site was prepared with 20 mL of 5% povidone-iodine solution (Zhejiang Apeloa Inc., China). A double-T incision was made in the nasal region, and the periosteum was carefully dissected to expose the nasal bone. The nasal bone was then removed to access the anterior bony wall of the sphenoid bone. The middle and posterior olfactory nerve filaments were excised using an endoscopic microdebrider to obtain a clear surgical field. An artificial sphenoid sinus was created to expose the chiasmatic optic nerve within the sphenoid bone. Using an endoscopic microdrill, circular windows were created at the center and on both sides of the optic nerve canal (chiasmatic and bilateral intracanalicular segments) to further expose the optic nerve.

### Retrobulbar Optic Nerve Exposure and Electrode Implantation in Goats

The procedure for lateral orbitotomy to expose the retrobulbar optic nerve was adapted from a previously described method*. Briefly, following exposure of the optic chiasm, a 5-6 cm continuous incision was made on the lateral side of the orbit. The orbital bone was partially removed after blunt dissection of the periosteum. During the exposure of the optic nerve, surgical sutures were used to retract the extraocular muscles, preventing potential damage. A needle electrode was then implanted into the retrobulbar segment of the optic nerve. The procedure was performed bilaterally to ensure symmetrical exposure and implantation.

### Retrobulbar Electrode Implantation in Rats

To expose the retrobulbar optic nerves of the rats, the instruments were sterilized, and the surgical area was prepared. The rats were anesthetized with isoflurane. A 5 mm incision was made over the lateral canthus, and the region was carefully dissected through the subcutaneous tissue to expose the underlying muscles. Blunt dissection was then performed to access the retrobulbar space, isolating the optic nerve from surrounding tissues. A cuff electrode measuring 2 mm in length with an internal diameter of 1 mm (slightly wider than the optic nerve of the rat) and a thickness of 0.3 mm was subsequently implanted on the optic nerve and securely fixed. The lateral rectus muscle was repositioned, and the skin incision was sutured with 6-0 nylon. The rat was monitored during recovery on a heating pad. Sham-operated group underwent the same surgical exposure of retrobulbar optic nerve but did not receive a cuff electrode implantation. Both the experimental group and the sham-operated group of rats were fitted with Elizabethan collars to prevent them from scratching the wounds and electrodes, which could affect the experimental results.

### Quantification of RGCs in Rats

The rodents were euthanized with an overdose of isoflurane, and their enucleated eyes were dissected to form posterior segment eyecups, which were then fixed in 4% paraformaldehyde for 2 hours at room temperature. The eyecups were subsequently dissected to create retinal flat mounts. The flat-mounted retinas were blocked for 2 hours at room temperature with blocking buffer containing 10% normal donkey serum and 0.3% Triton X-100 in PBS. The primary antibody (RBPMS; 1:400; Proteintech, 15187-1-AP) was diluted in blocking buffer and incubated for 24 hours at 4 °C on a shaker. After the flat mounts were washed in PBS with 0.3% Triton five times for 15 minutes each, they were incubated overnight with Alexa Fluor 488-conjugated anti-rabbit secondary antibodies. Following additional washes with PBS, the stained retinas were imaged using confocal microscopy (Cell Observer SD, Zeiss, Germany). Laser wavelength: 488nm; gain: 1000; exposure time: 50ms. To analyze the RGC density, multiplane z-series images were collected with a 20× objective, spanning from the nerve fiber layer (NFL) to the retinal pigment epithelium (RPE). The RGC density was calculated by imaging 12 retinal sections from the central to the peripheral regions across all four quadrants and averaging the values.

### Statistical Analysis

Statistical analyses were conducted using GraphPad (9.0) software. Correlation analysis was performed between the measured and simulated data. Two-way ANOVA was used to analyze potential differences and RGC data. Statistically significant differences are indicated by asterisks (* p<0.05, ** p<0.01, *** p<0.001, **** p<0.0001). The data are presented as the means ± s.e.m.s.

## Author contributions

Shengjian Lu: Data curation, Formal analysis, Software, Validation, Visualization, Writing _–_ original draft, Writing _–_ review & editing; Tonghe Yang: Investigation, Software, Validation, Visualization, Writing _–_ original draft; Gengyuan: Investigation, Validation, Writing _–_ review & editing; Huan Wu: Investigation, Methodology; Yangrui Huang: Investigation, Methodology; Te Zheng: Investigation, Methodology; Haodi Chen:Visualization; Shurui Huang: Writing _–_ review & editing; Yi Cao: Writing _–_ review & editing; Jian Yang: Writing _–_ review & editing; Wentao Yan: Funding acquisition, Writing _–_ review & editing; Yikui Zhang: Conceptualization, Data curation, Formal analysis, Funding acquisition, Project administration, Supervision, Writing _–_ original draft, Writing _–_ review & editing; Wencan Wu: Funding acquisition, Project administration, Supervision.

## Data availability statement

Requests for information and resources should be directed to the lead contact, Yikui Zhang. The data of the head model is available on GitHub at https://github.com/LuuSJ/Head-Model-for-Optic-Nerve-BMI. (accessible after accepted)

## Acknowledge

The research was funded by the following foundations:

National Key R&D Program of China (2022YFA1105501)

National Natural Science Foundation of China (82471080; 82171048; 81800842)

Key R&D Program of Zhejiang Province (2021C03065)

Key R&D Program of Wenzhou Eye Hospital (YNZD1201902)

Key R&D Program of Wenzhou (ZY2022021)

Key R&D Program of Wenzhou (H20220008)

## Ethics approval statement and patient consent statement

Ethical approval was granted by the Ethics Committee of the Eye Hospital of Wenzhou Medical University (Approval No. 2024-204-K-171-01), and informed consent was obtained from each volunteer. The animal protocols were approved by the Experimental Animal Ethics Committee of Wenzhou Medical University (Approval No. wydw2024-0241).

## Supplemental information

**Figure S1.**
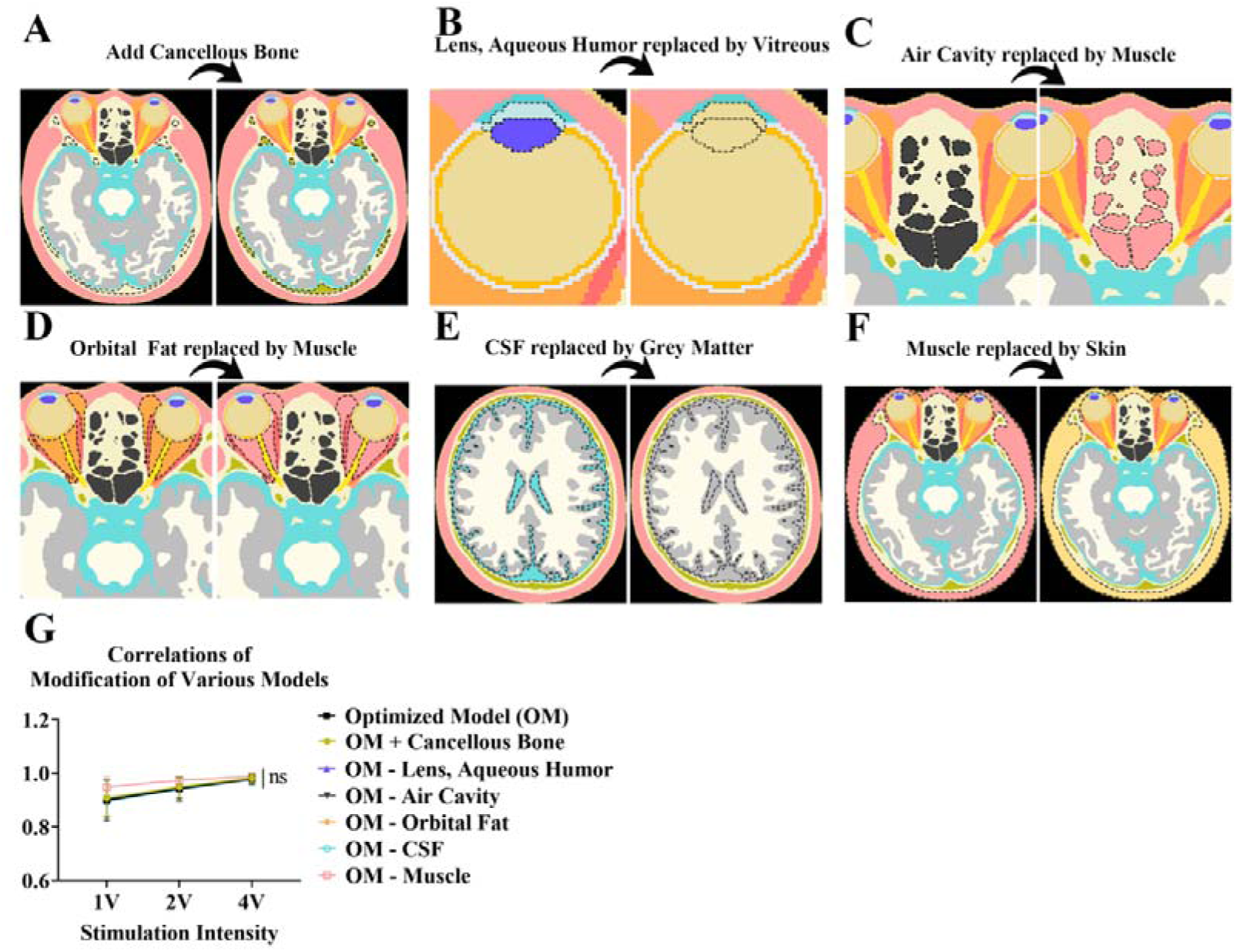
Impact of different model components on the accuracy of the human models. Illustrations of various modifications to the model components. **(A)** Addition of cancellous bone. **(B)** Replacement of the lens and aqueous humor with vitreous humor. **(C, D)** Replacement of the air cavities and orbital fat with muscle. **(E)** Replacement of the CSF with gray matter. **(F)** Replacement of the muscle tissue with skin. **(G)** Correlation analysis between simulated and measured data across three human volunteers with various stimulation intensities (1 V, 2 V, and 4 V) and different models.Two-Way Anova, n = 3, data are presented as mean ± s.e.m, ns: not significant.

**Figure S2.**
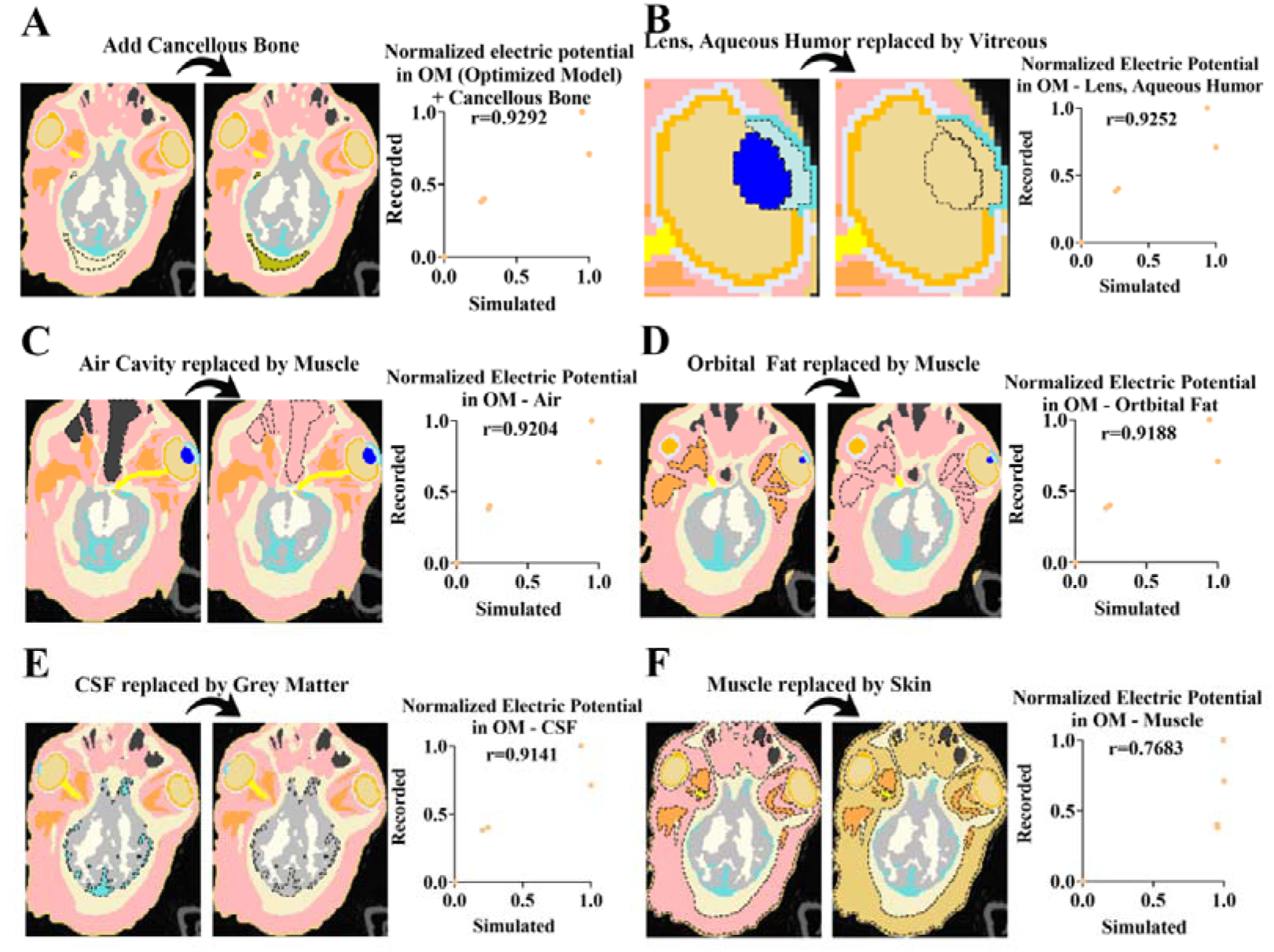
Impact of different model components on model accuracy for goat experiments. Illustrations of various modifications to the model components and the correlations between the normalized simulated and recorded electric potentials. **(A)** Addition of cancellous bone, r = 0.9292. **(B)** Replacement of the lens and aqueous humor with vitreous humor, r = 0.9252. **(C)** Replacement of air cavities with muscle, r = 0.9204. **(D)** Replacement of orbital fat with muscle, r = 0.9188. **(E)** Replacement of cerebrospinal fluid (CSF) with gray matter, r = 0.9141. **(F)** Replacement of muscle tissue with skin, r = 0.7683.

**Figure S3.**
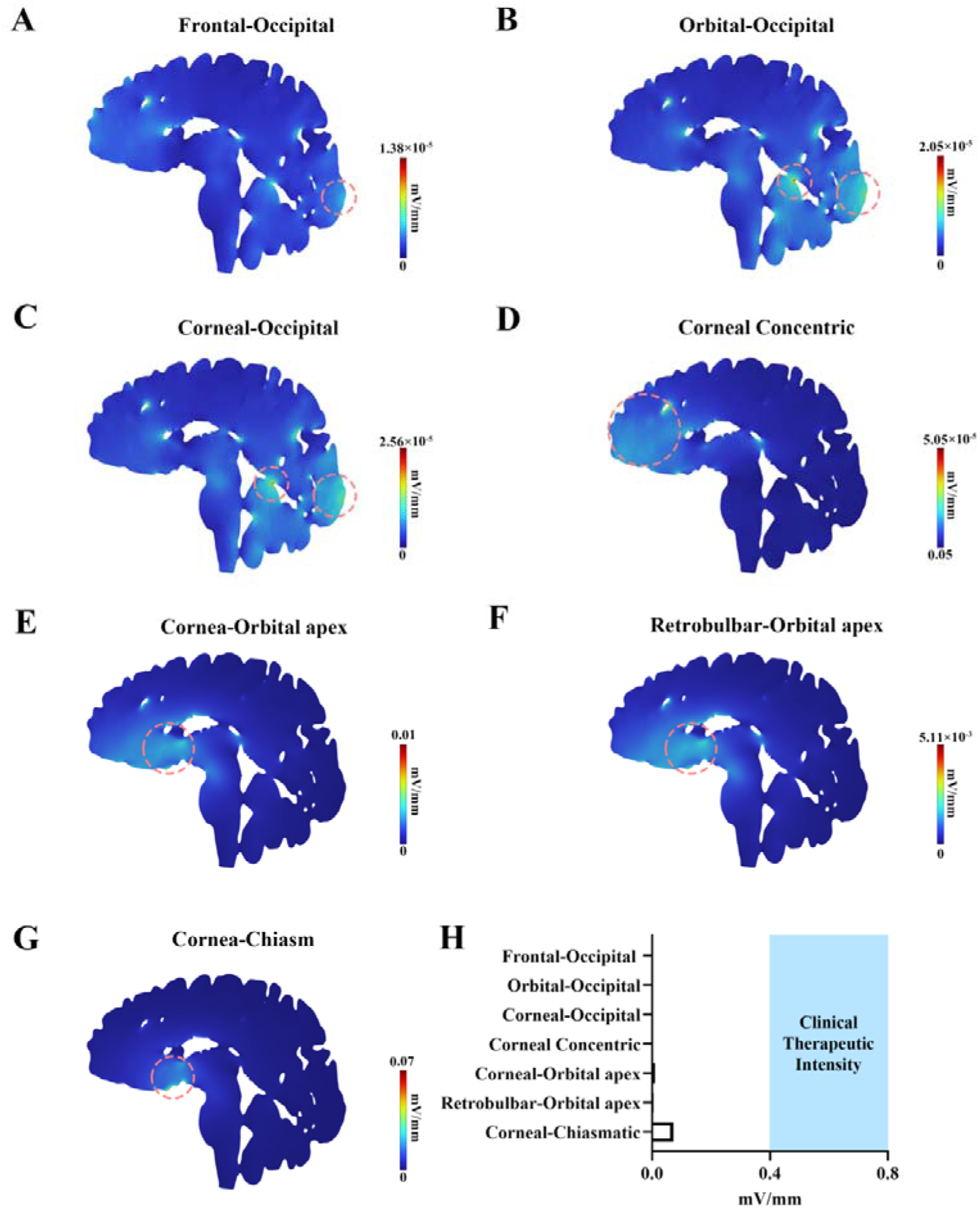
Distribution of the electric field strength in the sagittal brain regions of subject 1 across seven electrode placement strategies. Sagittal-view simulations of the electric field strength across different brain regions in volunteer 1 under seven electrode placement strategies: **(A)** frontal-occipital, **(B)** orbital-occipital, **(C)** corneal-occipital, **(D)** corneal concentric, **(E)** corneal-orbital apex, **(F)** retrobulbar-orbital apex, and **(G)** corneal-chiasmatic.

**Figure S4.**
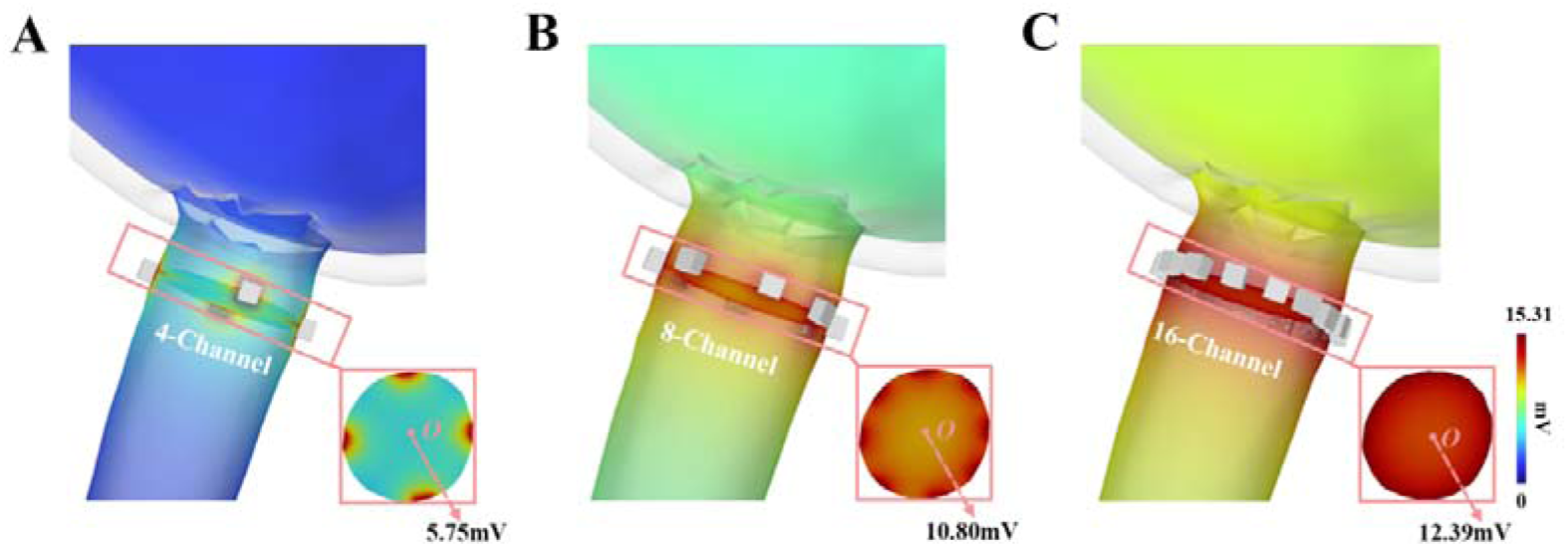
Electric potential distribution of various channel configurations. **(A-C)** Schematics of 4 to 16-channel configurations and the cross-sections of the electrode sites.

